# Early-life immune expression profiles predict later life health and fitness in a wild rodent

**DOI:** 10.1101/2021.10.08.463659

**Authors:** Klara M Wanelik, Mike Begon, Janette E Bradley, Ida M Friberg, Christopher H Taylor, Joseph A Jackson, Steve Paterson

## Abstract

Individuals differ in the nature of the immune responses they produce, affecting disease susceptibility and ultimately health and fitness. These differences have been hypothesised to have an origin in events experienced early in life that then affect trajectories of immune development and responsiveness. Here we investigate early life influences on immune expression profiles using a natural population of field voles, *Microtus agrestis*, in which we are able to monitor variation between and within individuals though time by repeat (longitudinal) sampling of individually marked animals. We analysed the co-expression of 20 immune genes in early life to create a correlational network consisting of three main clusters, one of which (containing *Gata3, Il10* and *Il17*) was associated with later life reproductive success and susceptibility to chronic bacterial (*Bartonella*) infection. More detailed analyses supported associations between early life expression of *Il17* and reproductive success later in life, and of increased *Il10* expression early in life and later infection with *Bartonella*. We also found significant association between an *Il17* genotype and the early life expression of *Il10*. Our results demonstrate that immune expression profiles can be manifested during early life with effects that persist through adulthood and that shape the variability among individuals in susceptibility to infection and fitness widely seen in natural populations.

## Introduction

Differences at birth, or experienced during neonatal or juvenile development, may impact an individual’s ability as an adult to respond to environmental allergens or pathogens, or to maintain health and reproduction (measures of which we define below). In medical sciences, the developmental origin of health and disease (DOMD) theory – which proposes that adult diseases can be traced back to childhood – has been highly influential since its inception and has provided insight into the aetiology of chronic inflammatory and allergic disorders in humans, such as atopy, asthma and autoimmune diseases^1,2^. Similarly, the development of the immune system during early life is an important influence on subsequent health and fitness^3^. The immune system is shaped by environmental exposure to allergens and pathogens, which, in turn, affect the immune responses made following subsequent exposure. Coupled with genetic polymorphisms affecting immune function^4^, early life effects associated with the immune system can give rise to substantial variation among individuals in their immune responses. Such environmental effects can act very early in life, even at birth. For example, children born by caesarean section are exposed to different microbes to vaginal delivery. This has been shown to modify their gut microbiota and the production of pro-inflammatory factors, and ultimately to increase their risk of opportunistic infections^5^ and chronic inflammatory disorders in later life e.g. asthma, type 1 diabetes and celiac disease^6,7^.

While effective immune responses are essential to health by protecting individuals against pathogens, immunopathology (such as tissue damage from inflammatory or over-reactive responses) contributes to significant ill-health in chronic inflammatory and allergic disorders such as atopy, asthma, autoimmune diseases or indeed infectious diseases such as COVID-19. An important concept is the hygiene hypothesis, which proposes that microbial exposure early in life helps the development of a well-adjusted immune system, the lack of which during childhood in modern societies contributes to the development of immunopathology and chronic immune disorders. Many studies have shown that traditional farming methods where children are widely exposed to microbes from an early age, develop fewer chronic immune disorders than children brought up in modern farms where there are lower levels of microbial exposure. This has been referred to as the ‘farm effect’^8,9^. Variation in traits associated with immunopathology can also be found in natural populations. In a wild population of moose, malnutrition early in life is linked to both a greater risk of osteoarthritis later in life (a disease associated with immune dysregulation) and reduced life expectancy^10,11^. In a feral population of Soay sheep, heritable variation in levels of anti-nuclear antibodies – which are self-reactive and if maintained at high titres with other markers are indicative of immune dysregulation in humans and dogs – are associated with variations in reproduction and over-winter survivorship^12^. Indeed, natural populations can provide a useful model to understand early life effects on the immune system, because they consist of genetically diverse individuals living in an environment that naturally exposes their immune systems to a wide array of pathogens and allergens. The interaction of different immune components with these environmental stimuli may lead to differences among individuals in their capacity for either effective responses to infection or immunopathologies.

Previous studies of early-life effects on immunity in natural populations have considered few immune parameters. For example, a number of avian studies have shown associations between broad-brush measures of nestling immune function, such as responsiveness to phytohemagglutinin (PHA) injection, and recruitment^13–16^. However, immune responses involve a complex network of interacting signalling and effector molecules, controlling pathways specialised at combating different types of pathogens, or mitigating damage caused by the pathogen or by immunopathology. Activity in one pathway of the immune system may have either positive or negative effects on the activity of another immune pathway. Similarly, the activity of immune pathways will both be influenced by environmental cues, such as infection and nutrition, and, in turn, influence disease resistance, health and fitness. A more nuanced and sophisticated approach to study immunity in natural systems is therefore needed, which considers multi-dimensional immune phenotypes; providing a way to reduce the complexity of these phenotypes, while still acknowledging the links between immune pathways. The availability of genome and transcriptome sequences provides new possibilities to measure immune phenotypes in natural populations, particularly through reverse transcription quantitative PCR (Q-PCR) to measure transcriptional expression of key immune regulators such as cytokines that are indicative of activity within different immune pathways^17^.

Here, we investigate early life influences on immune expression profiles in a natural population of field voles, *Microtus agrestis*. In this study system we are able to sample individuals longitudinally through their lives to measure a range of immunological parameters using Q-PCR and to relate these to subsequent variations in measures of health and fitness, including responses to infection. We do so having initially sought structure in the patterns of expression in our immune network, allowing us to characterise immune expression profiles in terms of appropriate clusters of genes, rather than single markers. Our central aim is to understand the extent to which, (1) the reproductive success of mature individuals (a measure of fitness), and (2) the response to infection of mature individuals (a measure of health) is foreshadowed by immune expression in the early stages of their lives. We note at the outset that the answers to these three questions are likely to vary between different parts of the immune network, and hence a further aim of this study is to begin to elaborate the range of possible patterns, and not to establish a general rule.

## Methods

*M. agrestis* were live-trapped from natural populations in Kielder Forest, Northumberland, UK, from 2015-2017 across seven different sites, each a forest clear-cut. At each site, 150-197 Ugglan small mammal traps (Grahnab, Sweden) were laid out in a grid spaced approximately 3-5 m apart. Our study was divided into longitudinal (individuals sampled live, multiple times) and cross-sectional (individuals sampled following culling) components.

### Ethics statement

All animal procedures were performed with approval from the University of Liverpool Animal Welfare Committee and under a UK Home Office licence (PPL 70/8210 to S.P.).

For the longitudinal study, on first capture, voles were injected with a Passive Integrated Transponder (PIT) tag (AVID, UK) for unique identification. On first and subsequent captures, a blood sample was taken whenever possible from the tail tip for immune gene expression assays and parasite detection, and on first capture a tail sample was taken for genotyping for pedigree reconstruction (see below). Other basic information was recorded including the snout-vent length, weight and pelage (for aging). Individuals were sampled in this way between 1-13 times. Some of these tagged voles (*n* = 510) were culled later in life for more detailed measurements to be taken (e.g., lens weight for aging purposes; see below), as part of our cross-sectional study.

Of the voles in the longitudinal study, 624 were first caught prior to breeding (at 6 weeks or less). These included juveniles and subadults, aged on weight and pelage. Based on previous work on the same study population, voles weighing 11g or less were assumed to be juveniles aged 2.5 weeks old (mean of 2-3 weeks) and voles weighing more than 11g but no more than 15g were assumed to be juveniles aged 4.5 weeks (mean of 3-6 weeks). Those voles weighing more than 15g but aged as juvenile on pelage, and those voles aged as subadult on pelage were assumed to be 6 weeks old. Finally, those voles aged as juvenile on pelage but with no weight measurement were assumed to be 4 weeks old (mean of 2-6 weeks)^18,19^. Of these 624 voles known to be first caught prior to breeding, 195 had a blood sample taken at first capture.

Some individuals may have first been caught as subadults but not recorded as such due to their adult-like pelage. In order to identify these individuals (and to increase our sample size) we regressed age against eye lens weight for those voles definitely first sampled prior to breeding (for which we had a birth date) that were culled in later life as part of our cross-sectional study (*n* = 99). The relationship between age and eye lens weight has been previously described^20^. A quassipoisson GLM with quadratic and cubic terms for eye lens weight, and log link, provided a good model fit for our data (*r*^*2*^ = 0.56; Supplementary Fig. S1). We used this regression to predict the ages (and hence birth dates) for all other culled voles. Of these, 59 voles were inferred to have first been caught prior to breeding, since their birth date was no more than 6 weeks before their date of first capture, of which 28 had a blood sample taken at first capture. Therefore, the final number of voles first sampled prior to breeding (known and inferred) was 223.

### Immune gene expression assays

We used SYBR green based Q-PCR to measure the expression levels of a panel of 20 immune-associated genes (Table 1) in blood from our longitudinal animals. The choice of our panel of genes was informed by (1) known immune-associated functions in mice, combined with (2) significant sensitivity of gene expression to environmental or intrinsic host variables in our previous studies^21,22^ or in a recent differential expression analysis of RNASeq data (not reported here).

**Table 1.**
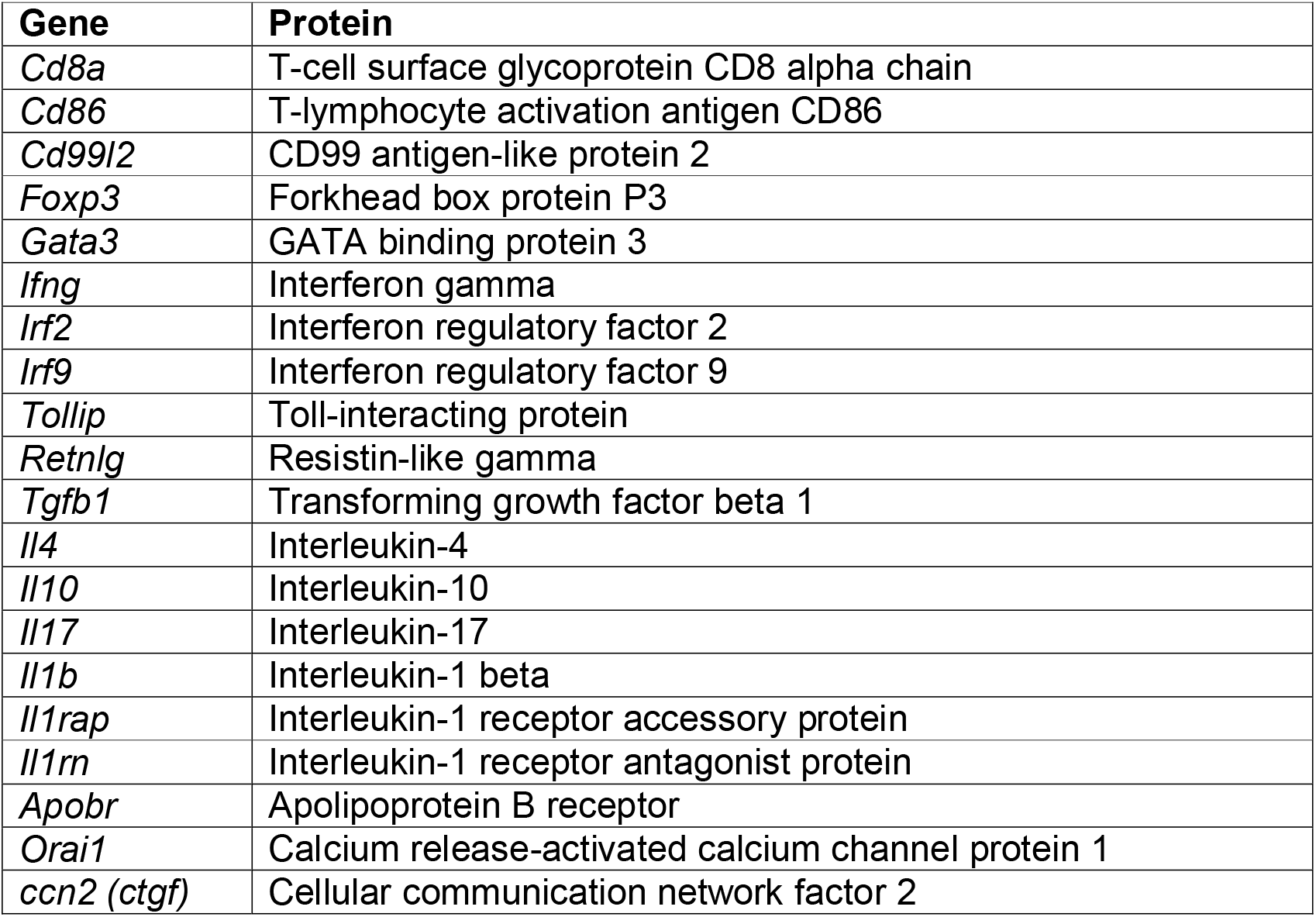
Panel of 20 immune-associated genes for which expression levels in blood samples were measured using Q-PCR.

All primer sets were designed *de novo* in-house and validated (to confirm specific amplification and 100±10% PCR efficiency under assay conditions). *Ywhaz* and *Actb* were employed as endogenous control genes. We extracted RNA from blood conserved in RNAlater using the Mouse RiboPure Blood RNA Isolation Kit (ThermoFisher), according to manufacturer’s instructions. RNA extracts were DNAse treated and converted to cDNA using the High-Capacity RNA-to-cDNA™ Kit (ThermoFisher), according to manufacturer’s instructions, including reverse transcription negative (RT-) controls for a subsample. SYBR green-based assays were pipetted onto 384 well plates by a robot (Pipetmax, Gilson) using a custom programme and run on a QuantStudio 6-flex Real-Time PCR System (ThermoFisher) at the machine manufacturers default real-time PCR cycling conditions. Reaction size was 10 µl, incorporating 1 µl of template and PrecisionFAST qPCR Master Mix with low ROX and SYBR green (PrimerDesign) and primers at the machine manufacturer’s recommended concentrations. We used three standard plate layouts for assaying, each of which contained a fixed set of target gene expression assays and the two endogenous control gene assays (the same sets of animals being assayed on matched triplets of the standard plate layouts). Unknown samples were assayed in duplicate wells and calibrator samples in triplicate wells and no template controls for each gene were included on each plate. Template cDNA (see above) was diluted 1/20 prior to assay. A main calibrator sample (identical on each plate) was created by pooling cDNA from blood samples taken from many different voles from the study site. As *Tollip, Il1rap* and *Irf2* were relatively poorly represented in this main calibrator sample, a synthesised 478 bp gene fragment containing the amplification target for each of these genes was used as an additional calibrator sample (at 10 × 10^5^ copies µl^-1^) in these cases. Samples from different field sampling groups were dispersed across plate triplets, avoiding confounding of plate with the sampling structure. Gene relative expression values used in analyses are RQ values calculated by the QuantStudio 6-flex machine software according to the ΔΔCt method, indexed to the appropriate calibrator samples. Melting curves and amplification plots were individually inspected for each well replicate to confirm specific amplification.

### Parasite detection

We quantified infections by microparasites (*Babesia microti* and *Bartonella* spp.) in blood using a strategy analogous to the host gene expression assays above, targeting pathogen ribosomal RNA gene expression and normalizing to the host endogenous control genes. We included two extra sets of primers in the blood Q-PCR assays described above (in one of the standard plate layouts for each plate triplet above). For *B. microti* we used the forward primer CTACGTCCCTGCCCTTTGTA and reverse primer CCACGTTTCTTGGTCCGAAT targeting the 18S ribosomal RNA gene and for *Bartonella* spp. we used the forward primer GATGAATGTTAGCCGTCGGG and reverse primer TCCCCAGGCGGAATGTTTAA targeting the 16S ribosomal RNA gene. As a calibrator sample we employed the main calibrator sample used for host gene expression in blood (above) in addition to a pool of DNA extracted from 154 blood samples from different *M. agrestis* at our study sites in 2015 and 2016; these DNA extractions were carried out using the QIAamp UCP DNA Micro Kit (Qiagen) following manufacturer’s instructions.

Relative expression values are presumed to relate to the expression of pathogen ribosomal RNA genes and in turn to the intensity of infection. Pilot testing indicated that the detected relative quantities of parasite RNA correlated closely to visual counts of parasitized cells on blood smears examined under a microscope. Moreover, we validated our diagnostic results by comparing our PCR RQ values to independent data for a subset of cross-sectional voles with mapped genus-level pathogen reads from RNASeq analysis of blood samples (*n* = 44), finding the two data sets strongly corroborated each other (*Bartonella* spp., *r*^*2*^ = 0.81, *p* < 0.001; *B. microti, r*^*2*^ = 0.81, *p* < 0.001).

### Genotyping & pedigree reconstruction

We genotyped voles from their tail samples for 346 single nucleotide polymorphisms (SNPs) in 127 genes. See Wanelik *et al*.^23^ for details of the approach used to select these SNPs. Briefly, we combined information from an external database of human immune genes (Immunome database; http://structure.bmc.lu.se/idbase/Immunome/index.php) with our own existing knowledge of the study system. DNA was extracted from a tail sample taken from the animal using DNeasy Blood and Tissue Kit (Qiagen). Genotyping was then performed by LGC Biosearch Technologies (Hoddesdon, UK; http://www.biosearchtech.com) using the KASP SNP genotyping system. This included negative controls (water) and duplicate samples for validation purposes.

We used a subset of our SNP dataset (*n* = 114 SNPs) to reconstruct a pedigree using the R package *Sequoia*^24^. All analyses were performed in R statistical software version 3.5.2^25^. Full details can be found in Wanelik *et al*.^26^. Briefly, we inputted life history information into *Sequoia* where possible (99% of samples were assigned a sex; 54% were assigned a birth month), and generated site-specific pedigrees (assuming no dispersal between sites, each several kilometres apart). We inspected log10 likelihood ratios (LLRs) for parent pairs as recommended in the user manual for *Sequoia*. Almost all LLRs were positive (97% of LLRs) indicating confidence in our assignments. For each individual present in a pedigree (*n* = 652; see below), the number of offspring was counted to provide a measure of their reproductive success. Half of individuals present in our pedigrees (*n* = 325) were found to have no offspring. We expect the majority of these to be true zeros (representing actual reproductive failure) as we sampled a large proportion of the total population within clear-cuts. We minimised the chance of false zeros by excluding from the pedigree those individuals (e.g., at the periphery of a study grid) for which we recorded no relatives (including offspring) likely because we had not sampled in the right place.

## Statistical methods

### Correlation networks

We constructed an expression correlation matrix for all immune genes measured prior to breeding, in early life. We used Spearman Rank correlation coefficients (1) in case of any non-linear relationship between genes, and (2) because our expression data were not normally distributed. There was some missing data in our expression dataset (640/4460 values = 14%). In order to maximise the sample size used to generate each correlation coefficient, we used pairwise complete observations. This resulted in 72% of correlations based on at least 75% of the total data. For each correlation coefficient, we randomly permuted the data 1000 times to calculate a *p*-value. We also adjusted all *p*-values for multiple testing using the Benjamini-Hochberg method. We thresholded our correlation network using these corrected *p*-values – only keeping those edges with a significant corrected *p*-value (*p* ≤ 0.05). Genes were clustered on edge betweenness, using the edge betweenness algorithm in igraph^27^. The edge betweenness score of an edge is a measure of the number of shortest paths that go through it. The algorithm identifies densely connected modules by gradually removing edges with highest edge betweenness scores^28^.

We then ran an exploratory analysis, repeating the steps above to construct a correlation network for all immune genes measured in early life and six additional measures of later life or lifetime success. These were: (1) whether or not an individual was recaptured (a proxy for survival), (2) reproductive success in later life (number of offspring in the pedigree), the proportion of later life infected with two major microparasites in our population, namely (3) the bacterium *Bartonella* spp. and (4) the protozoan *Babesia microti*, (5) mean Scaled Mass Index (SMI; a measure of an individual’s average condition across their lifetime^29^), and (6) coefficient of variation in SMI (a measure of an individual’s resilience in maintaining their condition across their lifetime; with the caveat that this measure of lifetime of success will be less reliable when based on fewer captures; see Supplementary Fig. 2). There was a lack of variation in the first of these measures, whether or not an individual was recaptured (total of 191 recaptured and 32 not recaptured) which led to zero variance warnings and missing correlation coefficients for some pairwise comparisons. This measure was therefore omitted from the network and instead tested for associations with immune parameters of interest using a Fisher’s exact test (see below). There were slightly more missing data in our final combined expression and success dataset (1078/5575 values = 19%). As before, we used pairwise complete observations. This time, with 45% of correlations based on at least 75% of the total data.

### Confirming associations between immune genes and measures of later life success

Follow-up analyses were run to confirm the two correlations of interest: (1) a significant positive correlation between *Il17* expression in early life and reproductive success, and (2) a significant positive correlation between *Il10* expression in early life and proportion of later life infected with *Bartonella* spp. (see Results).

We had information about both *Il17* expression in early life *and* reproductive success for 131 voles. Our measure of reproductive success was zero-inflated (107/131 or 82% zero values) and was simply coded as either reproduced or not. We then ran a logistic regression to model the probability of reproducing. *Il17* expression in early life was also found to be zero-inflated (117/131 or 89% zero values) and was simply coded as either expressed or not. We included birth month as a (continuous) covariate in the model, given that autumn-born voles have a lower chance of reproducing than spring-born voles^30^. Other (categorical) covariates included in the model as fixed effects were: sex, whether or not an individual was culled for the cross-sectional component of this study (again, reducing the opportunity to reproduce) and the year in which an individual was born. We also included a random effect for the site in which an individual was born.

We had information about *Il10* expression in early life *and* future *Bartonella* infection for 104 voles. *Bartonella* infection status was assessed multiple times for the majority of individuals in the longitudinal component of the study (mean = 2.3; range = 1-6). Therefore, we ran a logistic regression, with a random effect for individual, to model the probability of an individual being infected throughout their life. The response variable was whether (1) or not (0) an individual tested positive at a particular time. As was the case for *Il17, Il10* expression in early life was found to be zero-inflated (89/104 or 86% zero values) and was simply coded as either being expressed or not. Other individual-specific covariates, considered potential drivers of future *Bartonella* infection and included as fixed effects were: sex, whether or not an individual was targeted for anti-parasite treatment (which may have affected the flea vectors of *Bartonella*), whether or not they were already infected with *Bartonella* in early life and birth year. We also included a random effect for birth site.

Logistic regressions were run using the R package glmmADMB^31,32^. All covariates were tested for independence using variance inflation factors (all VIFs < 3). Full submodel sets were generated from each global model, including all fixed and random terms of interest, using the MuMIn package (Bartoń 2016). All candidate models were then evaluated and ranked on relative fit using the Akaike Information Criterion, AIC. The best model, with the lowest AIC, is reported in the text and figures.

### Testing the age-specificity of these associations

To test whether these associations are strictly early-life or also present in adult (breeding) samples, we re-ran the same best models for reproductive success and proportion of later life infected with *Bartonella* on adult blood samples. We maximised the sample size for these analyses by including all individuals sampled as adults, whether or not they were first sampled prior to breeding and therefore had an approximate birth date. For this reason, we were unable to account for birth month (reproductive success) and birth year (reproductive success and *Bartonella* infection). The best models were otherwise unchanged. After omitting those samples with missing data, we had information about both *Il17* expression in adult life *and* reproductive success for 357 voles, and information about both *Il10* expression in adult life and *Bartonella* infection for 798 voles. For individuals with more than one adult sample, we selected one adult sample at random from which to measure expression. To ensure that our results were robust to different random draws, we repeated this process 1000 times, re-running the best models on 1000 random draws. We report the number of random draws for which each association was significant (*p* ≤ 0.05) compared to that expected by chance. We also tested for an association between whether or not an individual expressed each immune parameter of interest early in life, and whether or not they expressed it in later life using a Fisher’s exact test.

### Testing for associations between immune genes of interest and genetic polymorphisms

We looked for associations between the immune parameters of interest (*Il10* and *Il17* expression in early life) and genetic polymorphism. In order to avoid difficulties in interpretation due to multiple testing, we looked only at polymorphisms in the immune genes found within immune cluster 3 (see Results and Fig. 1; *Il17* = 1 SNP, *Il10* = 2 SNPs, *Gata3* = 2 SNPs). We used the R package hapassoc (for *Il10* and *Gata3*, for which we had information about more than one SNP) and the package SNPassoc (for *Il17*, for which we had information about a single SNP). Both hapassoc and SNPassoc models assumed an additive genetic model.

**Fig. 1.**
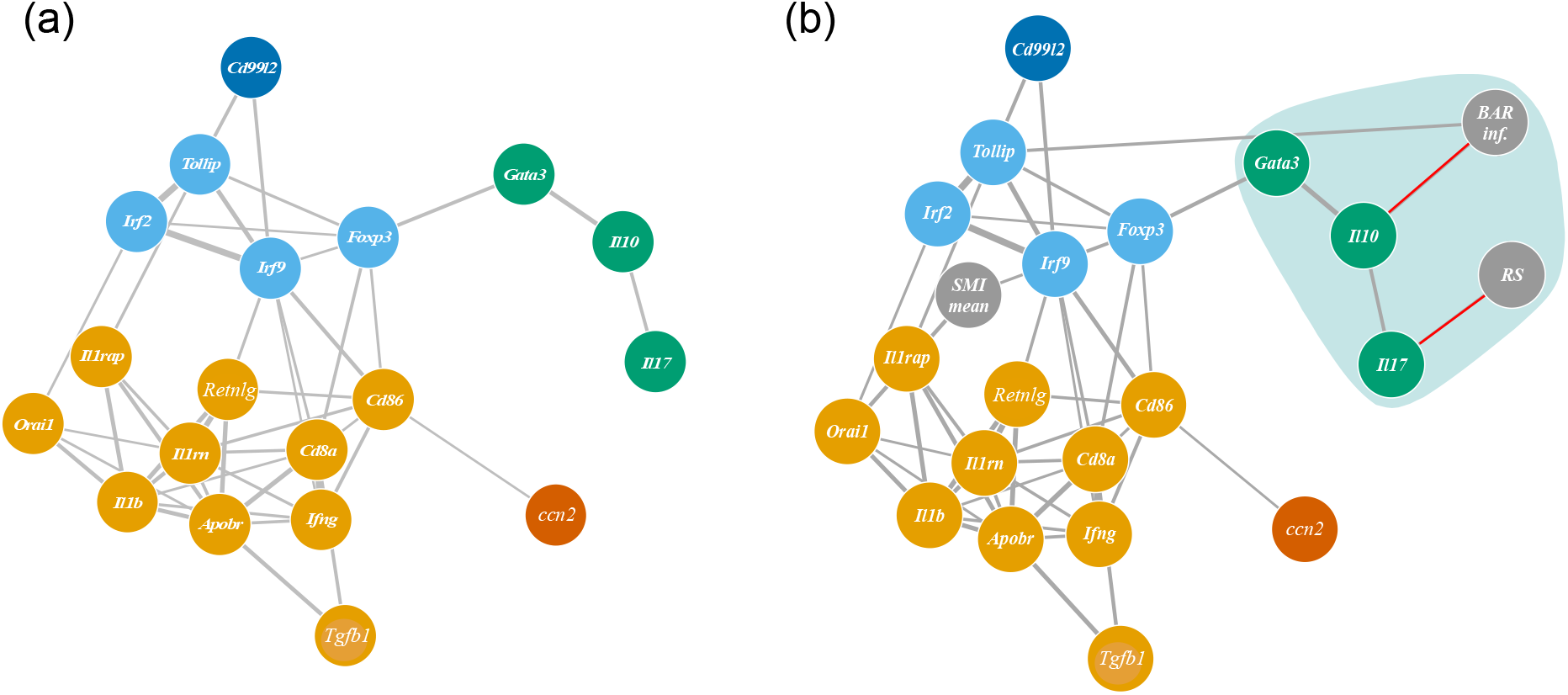
Network showing significant correlations (and associated nodes) of magnitude ≥ 0.2 between (a) immune parameters in early life (coloured nodes) and (b) immune parameters in early life and measures of future success (grey nodes). Nodes without any significant correlations of magnitude ≥ 0.2 are not shown in either (a) or (b). Width of an edge represents exact magnitude of the Spearman rank correlation coefficient. Three main clusters of immune parameters emerge based on edge betweenness (indicated by the colour of nodes in (a): orange, light blue and green). One of these (cluster with green nodes and also highlighted in green) was closely associated with two measures of future success (*Bartonella* infection and reproductive success) and was investigated further, in particular the edges highlighted in red in (b).

## Results

### Three main clusters of immune genes are visible in early life

We constructed a correlation network for 20 immune genes whose expression was measured in our voles prior to breeding. Following thresholding, one immune gene, *Il4*, dropping out of the network. Using clustering analysis we identified three main immune clusters in early life: (1) immune cluster 1 (*Il1rap, Il1rn, Il1b, Orai1, Retnlg, Apobr, Cd8a, Ifng, Tgfb1, Cd86*), (2) immune cluster 2 (*Irf2, Irf9, Foxp3, Tollip*) and (3) immune cluster 3 (*Gata3, Il10, Il17*; Fig. 1a).

### Exploratory analysis points to immune cluster 3 being closely associated with measures of later life success

We then ran an exploratory analysis, repeating the same process as above, but adding six measures of later life success. Two measures of later life success (coefficient of variation in SMI and proportion of later life infected with *Babesia*) were not correlated with any other measure and dropped out of the network. Both immune clusters 1 and 2 were associated with mean SMI. This included a significant negative correlation between *Orai1* expression in early life (immune cluster 1) and mean SMI (*rho* = -0.33; corrected *p* < 0.01), and a significant positive correlation between *Foxp3* expression in early life (immune cluster 2) and mean SMI (*rho* = 0.23; corrected *p* < 0.05). However, the immune cluster with the most associations with measures of later life success was immune cluster 3. Two measures of later life success (reproductive success and the proportion of later life infected with *Bartonella* spp.) were associated with immune cluster 3 (cluster highlighted in green in Fig. 1b). This included a significant positive correlation between *Il17* expression in early life and reproductive success (*rho* = 0.22; corrected *p* = 0.04), and a significant positive correlation between *Il10* expression in early life and proportion of later life infected with *Bartonella* spp. (*rho* = 0.22; corrected *p* = 0.05; edges highlighted in red in Fig. 1b).

### Follow-up analysis confirms that *Il17* expression in early life is associated with increased odds of reproducing later in life

A logistic regression confirmed that whether or not an individual expressed *Il17* in early-life was significantly associated with their probability of reproducing in later life (odds ratio = 5.10; 95% CI = 1.28 – 20.32; *p* = 0.02). Other fixed effects which appeared in the best model for, and were significantly associated with, the probability of reproducing in later life were the year in which an individual was born, and the month in which it was born (Fig. 2a). There was no significant association between *Il17* expression in later (breeding) life and reproductive success (*n* = 7/1000 random samples for which p ≤ 0.05; many fewer than the 50/1000 expected by chance) indicating that this association was specific to early-life expression of *Il17*. Consistent with this result, we found no association between whether or not an individual expressed *Il17* early in life, and whether or not they expressed *Il17* later in life (Fisher’s exact test; odds ratio = 0.64; 95% CI = 0.18 – 2.03; *p* = 0.45).

**Fig. 2.**
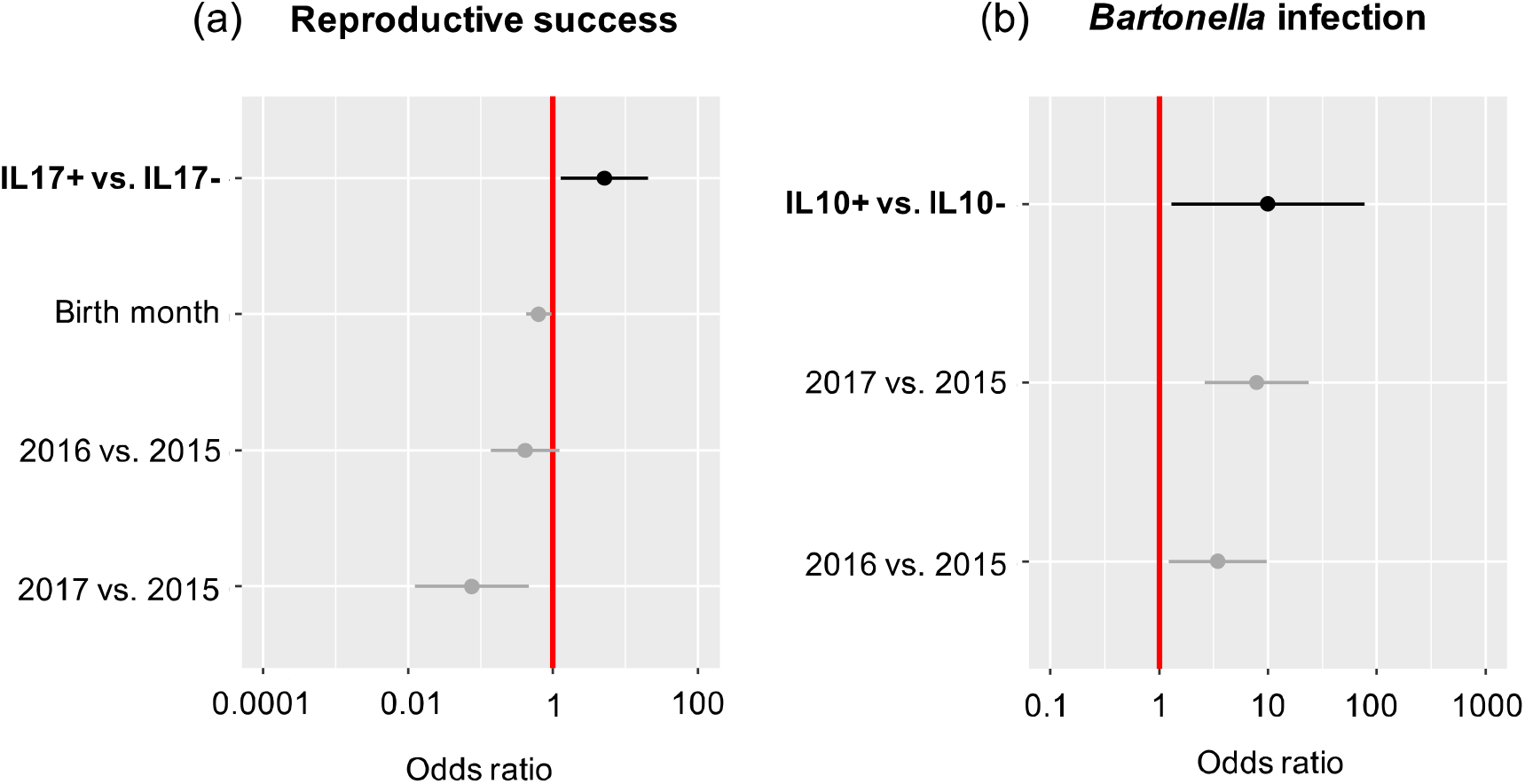
Odds of later-life success given immune parameters in early-life. In bold: (a) odds of reproduction for individuals expressing *Il17* in early life (IL17+) compared to individuals not expressing *Il17* (IL17-), and (b) odds of *Bartonella* infection for individuals expressing *Il10* in early life (IL10+) compared to individuals not expressing *Il10* (IL10-). Not in bold: odds of reproduction or infection associated with other fixed effects which also appeared in best models: year in which an individual was born (2015, 2016 or 2017) and month in which an individual was born (treated as continuous variable). All estimates taken from best models (see text). Error bars represent 95% confidence intervals.

### *Il10* expression in early life is associated with an increased proportion of later life infected with *Bartonella* spp. and decreased odds of recapture (a proxy for survival)

A logistic regression confirmed that whether or not an individual expressed *Il10* in early-life was significantly associated with their probability of being infected with *Bartonella* in later life (odds ratio = 9.98; 95% CI = 1.29 – 77.08; *p* = 0.003). Year in which an individual was born was the only other fixed effect present in the best model for, and was also significantly associated with, *Bartonella* infection (Fig. 2b). The association between *Il10* expression in later life and probability of infection was also significant (*n* = 993/1000 random samples for which p ≤ 0.05; many more than the 50/1000 expected by chance) indicating that this association was not specific to early-life expression. We found no association between whether or not an individual expressed *Il10* early in life, and whether or not they expressed *Il10* later in life (Fisher’s exact test; odds ratio = 1.05; 95% CI = 0.25 – 3.90; *p* = 1.00). *Bartonella* infection status in early life did not appear in the best model for explaining the probability of infection in later life, but there was a significant association between this variable and *Il10* expression in early life (Fisher’s exact test; odds ratio = 5.90; 95% CI = 1.38 – 53. 41; *p* = 0.01). In addition, we found a significant negative association between *Il10* expression in early life and probability of recapture (a proxy for survival; Fisher’s exact test, odds ratio = 0.39; 95% CI = 0.15 – 1.03; *p* = 0.05).

### *Il10* expression in early life is associated with a single SNP in the *Il17* gene

We looked for associations between the immune parameters of interest (*Il10* and *Il17* expression in early life) and genetic polymorphism. We found a significant association between a single SNP in the *Il17* gene and *Il10* expression in early life under an additive model, with the probability of expressing *Il10* significantly increasing with the number of T alleles (odds ratio = 1.74; 95% CI = 1.03 – 2.94; *p* = 0.04; Fig. 3). *Il17* genotype only explained a small proportion of variation in *Il10* expression (*r*^*2*^ = 0.026).

**Fig. 3.**
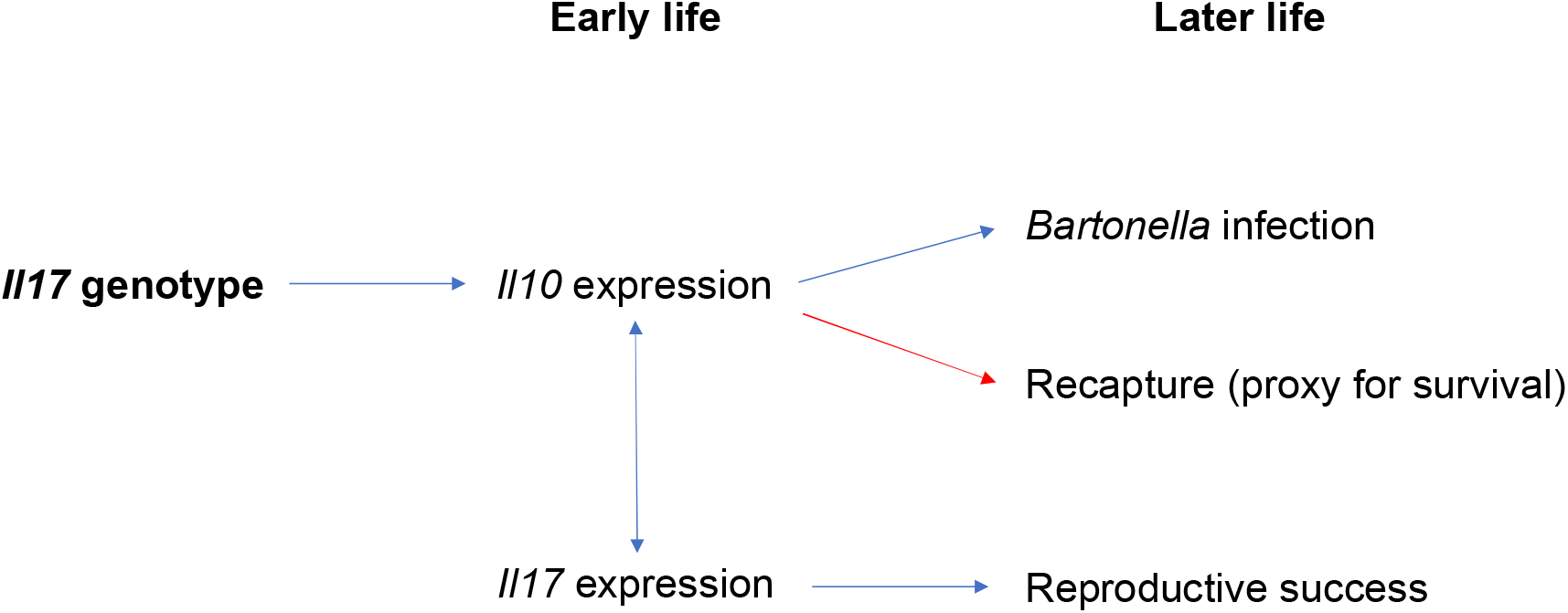
Summary diagram of confirmed associations. Blue arrows indicate positive associations, red arrows indicate negative associations. One-way arrows indicate a likely causal association, two-way arrows indicate a correlation only.

## Discussion

We have used a wild rodent system to provide evidence that, (1) the reproductive success (a measure of fitness), and (2) the response to infection of mature individuals (a measure of health) is foreshadowed by immune expression in the early stages of their lives. We have also found evidence to suggest that these patterns vary between different parts of the immune network.

We were able to derive three clusters of genes related to immune function based on co-expression data and it was unsurprising to see genes involved in different pathways, and with mixed pro- and anti-inflammatory functions, included in the same clusters. This is expected given the complexity of immune regulatory networks. For example, some responses might increase at the same time as their negative feedbacks, or responses within one pathway might lead to host states that favour the expression of another pathway. Nonetheless, we saw very clear functional biases, with the largest cluster containing predominantly pro-inflammatory markers such as *Il1* signalling cascade members^33^, *Ifng*^34^, *Tgfb1*^35^ and *Retnlg*^36^, but also genes with more nuanced functions in the promotion and suppression of inflammation, including *Tgfb1*^37^ itself and *Orai1*^38^. This “pro-inflammatory” cluster was distinct from a second cluster containing predominantly immune down-regulatory markers including *Foxp3*^39^, *Tollip*^40^ and *Irf2*^41^, but also I*rf9*^42^ a driver of interferon effector responses. A third cluster containing *Gata3, Il10* and *Il17* may give new insights into immune function, given that it contained a counterintuitive set of genes and was, at the same time, the cluster best linked to host health and fitness. We have previously found GATA3 to be associated with immune tolerance phenotypes, and the association with the anti-inflammatory cytokine IL10 provides further support of this finding^43,44^. *Il10* expression is, in turn, positively associated with *Il17* expression. IL17, which we found to be an early-life indicator of reproductive success, is typically associated with a pro-inflammatory immunopathology but may also be protective against the numerous primary microbial insults that young voles, lacking acquired immunity, may encounter^45^. The correlation between IL10 and IL17 may thus be one whereby IL10 is produced in response to IL17 in order to safely regulate its inflammatory effects and prevent immunopathology^46–49^. We argue that wild rodent studies can help us to define which of the possible immunological interactions known from laboratory studies predominate and shape the structure of immune phenotypes in individuals exposed to pathogens, nutritional challenge or other stressors within the natural environment. As in the case of IL10 and IL17, wild rodents may also give insights into immune regulation within a natural context.

Having identified the correlational structure of immunological traits in a wild population, it is then a natural extension to include other phenotypic traits. We used longitudinal data to address our aim to determine how immune phenotypes may be laid down in early life to affect later life health and fitness. We found two pathways through which immune phenotypes in early life act on later life traits (Fig. 3). In the first, increased expression of *Il10* in blood samples taken during early life (i.e., prior to breeding) was subsequently associated with both a ten-fold increased susceptibility to *Bartonella* infection later in life and a three-fold reduction in survivorship. The association of a genotypic effect on *Il10* expression, a polymorphism in *Il17*, suggests that early-life effects may be, at least in part, laid down at birth in this case. A challenge for future work will be to tease out causation from correlation in this case. An adverse *Il17* genotype may make an individual more likely to become infected with *Bartonella*, with infection stimulating *Il10* expression. Alternatively, that genotype may predispose an individual to express *Il10* early in life (and perhaps lead to other immune-expression variations) with that pattern of immune expression increasing the likelihood of *Bartonella* infection. The positive association between *Il10* expression in the earliest samples and *Bartonella* infection at that time supports but cannot positively establish the former, since *Bartonella* infection before even the earliest samples is possible. Further, it is unclear whether it is *Bartonella* infection or the pattern of immune expression that reduces survival.

We also found a significant association between I*l10* expression in breeding adults and the proportion of later life infected with *Bartonella*. It is perhaps not surprising that we found this association, as both *Il10* expression and *Bartonella* infection in this case were measured in contemporaneous samples (taken in later life) and our previous work shows that *Il10* expression increases soon after acquisition of *Bartonella*^50^. Furthermore, we found no significant association between *Il10* expression early and later in life, suggesting that the later-life association was not driven by its early-life counterpart.

The second pattern that we found is for increased *Il17* expression early in life to be associated with a subsequent five-fold increased likelihood of reproduction. By contrast, *Il17* expression in breeding adults showed no such association with reproduction, and indeed, no association either between *Il17* expression early and later in life. In contrast to *Il10* expression, no genetic effects on *Il17* expression were detected; at least from the deliberately limited set of cytokine polymorphisms considered, which included *Il17* polymorphism itself. This suggests that early life expression of *Il17* may be the consequence of an environmental effect leading to an immune phenotype that acts on juveniles and influences their development as they transition into reproductively mature adults. This could be, for example, a bacterial infection or composition of the gut microbiome early in life.

Hence, despite the contrasts between the patterns for *Il10* and *Il17* expression, both appear to be examples where the die has been cast early: immune expression early in life, however determined, is a powerful predictor of the future life course of the individual concerned. Previous evidence for this has been scarce. A number of studies have shown an association between broad-brush measures of immune expression in nestling birds and recruitment from the nest^13–16^, an almost contemporaneous effect. Bowers *et al*.^13^ went a step further, and showed an association between a broad-brush measure of immune expression (PHA responsiveness), longevity and reproductive success. Here we provide a more refined analysis, and describe two examples in which differences detectable early in life in the expression of distinct immune pathways play out much later in life, or at least over a much more extended period. *Il10* differences confirm a genetic basis for immune expression that, as previously shown in this system and others, act through the network of interacting cytokines such that polymorphism at one cytokine gene can affect expression of other cytokines and immune phenotypes^23,51–56^, with these effects further percolating to affect health and fitness more broadly in later life. By contrast, the *Il17* differences appear to be more reminiscent of the Jesuit motto “Give me the boy until he is seven and I will give you the man” : that independently of any genetic differences, an individual’s experiences early in life may, in this case through the medium of immune expression, profoundly influence their later health and fitness.

## Acknowledgements

The authors wish to thank those involved in obtaining and processing samples from the field: Rebecca Turner, Lukasz Lukomski, Stephen Price, Sarah Gore, Ed Parker, Maria Capstick, Noelia Dominguez Alvarez, Susan Withenshaw, William Foster, Ann Lowe, Benoit Poulin, Anna Thomason and Elena Arriero. They also wish to thank the Forestry Commission for access to the study sites and the Centre for Genomic Research at the University of Liverpool for sequencing samples. This research was funded by the Natural Environment Research Council (NERC) award NE/L013452/1 to S.P., M.B., J.A.J. and J.E.B.

## References

1. Barker, D. J. P. The origins of the developmental origins theory. J. Intern. Med. 261, 412–417 (2007).

2. Wadhwa, P., Buss, C., Entringer, S. & Swanson, J. Developmental Origins of Health and Disease: Brief History of the Approach and Current Focus on Epigenetic Mechanisms. Semin. Reprod. Med. 27, 358–368 (2009).

3. Dowling, D. J. & Levy, O. Ontogeny of early life immunity. Trends Immunol. 35, 299–310 (2014).

4. Brodin, P. et al. Variation in the human immune system is largely driven by non-heritable influences. Cell 160, 37–47 (2015).

5. Stewart, C. J. et al. Temporal development of the gut microbiome in early childhood from the TEDDY study. Nature 562, 583–588 (2018).

6. Khafipour, E. & Ghia, J. Mode of Delivery and Inflammatory Disorders. J. Immunol. Clin. Res. 1, 1004 (2013).

7. Shao, Y. et al. Stunted microbiota and opportunistic pathogen colonization in caesarean-section birth. Nature 574, 117–121 (2019).

8. Ober, C., Sperline, A., e, M. & Vercelli, D. Immune Development and Environment: Lessons from Amish and Hutterite Children. Curr. Opin. Immunol. 48, 51–60 (2017).

9. Von Mutius, E. & Vercelli, D. Farm living: Effects on childhood asthma and allergy. Nat. Rev. Immunol. 10, 861–868 (2010).

10. Peterson, R. O., Vucetich, J. A., Fenton, G., Drummer, T. D. & Larsen, C. S. Ecology of arthritis. Ecol. Lett. 13, 1124–1128 (2010).

11. Woodell-May, J. E. & Sommerfeld, S. D. Role of Inflammation and the Immune System in the Progression of Osteoarthritis. J. Orthop. Res. 38, 253–257 (2020).

12. Graham, A. L. et al. Fitness correlates of heritable variation in antibody responsiveness in a wild mammal. Science 330, 662–665 (2010).

13. Bowers, E. K. et al. Neonatal body condition, immune responsiveness, and hematocrit predict longevity in a wild bird population. Ecology 95, 3027–3034 (2014).

14. Cichon, M. & Dubiec, A. Cell-mediated immunity predicts the probability of local recruitment in nestling blue tits. J. Evol. Biol. 18, 962–966 (2005).

15. López-Rull, I., Celis, P., Salaberria, C., Puerta, M. & Gil, D. Post-fledging recruitment in relation to nestling plasma testosterone and immunocompetence in the spotless starling. Funct. Ecol. 25, 500–508 (2011).

16. Moreno, J. et al. Nestling cell-mediated immune response, body mass and hatching date as predictors of local recruitment in the pied flycatcher Ficedula hypoleucai. J. Avian Biol. 36, 251–260 (2005).

17. Turner, A. K. & Paterson, S. Wild rodents as a model to discover genes and pathways underlying natural variation in infectious disease susceptibility. Parasite Immunol. 35, 386–395 (2013).

18. Begon, M. et al. Effects of abundance on infection in natural populations: field voles and cowpox virus. Epidemics 1, 35–46 (2009).

19. Gebert, S. Patterns in helminth infections in natural populations of rodents. (University of Liverpool, 2008).

20. Rowe, F., Bradfield, A., Quy, R. & Swinney, T. Relationship Between Eye Lens Weight and Age in the Wild House Mouse (Mus musculusi). J. Anim. Ecol. 22, 55–61 (1985).

21. Jackson, J. A. et al. The analysis of immunological profiles in wild animals: A case study on immunodynamics in the field vole, Microtus agrestis. Mol. Ecol. 20, 893–909 (2011).

22. Jackson, J. A. et al. An immunological marker of tolerance to infection in wild rodents. PLoS Biol. 12, e1001901 (2014).

23. Wanelik, K. M. et al. A candidate tolerance gene identified in a natural population of field voles (Microtus agrestis). Mol. Ecol. 27, 1044–1052 (2018).

24. Huisman, J. Pedigree reconstruction from SNP data: parentage assignment, sibship clustering and beyond. Mol. Ecol. Resour. 17, 1009–1024 (2017).

25. R Core Team. R: A language and environment for statistical computing. R Foundation for Statistical Computing, Vienna, Austria. URL https://www.R-project.org/. (2018).

26. Wanelik, K. et al. IgE receptor polymorphism predicts divergent, sex-specific inflammatory modes and fitness costs in a wild rodent. biorXiv (2019).

27. Csardi, G. & Nepusz, T. The igraph software package for complex network research. InterJournal, Complex Syst. 1695 (2006).

28. Newman, M. E. J. & Girvan, M. Finding and evaluating community structure in networks. Phys. Rev. E 69, 026113 (2004).

29. Peig, J. & Green, A. J. New perspectives for estimating body condition from mass/length data: The scaled mass index as an alternative method. Oikos 118, 1883–1891 (2009).

30. Wang, M., Ebeling, M. & Hahne, J. Relevance of body weight effects for the population development of common voles and its significance in regulatory risk assessment of pesticides in the European Union. Environ. Sci. Eur. 31, (2019).

31. Fournier, D. A. et al. AD Model Builder: Using automatic differentiation for statistical inference of highly parameterized complex nonlinear models. Optim. Methods Softw. 27, 233–249 (2012).

32. Skaug, H., Fournier, D., Bolker, B., Magnusson, A. & Nielsen, A. Generalized Linear Mixed Models using ‘AD Model Builder’. R package version 0.8.3.3. (2016).

33. Dinarello, C. A. Overview of the IL-1 family in innate inflammation and acquired immunity. Immunological Reviews 281, 8–27 (2018).

34. Schroder, K., Hertzog, P. J., Ravasi, T. & Hume, D. A. Interferon-γ: an overview of signals, mechanisms and functions. J. Leukoc. Biol. 75, 163–189 (2004).

35. Han, G., Li, F., Singh, T. P., Wolf, P. & Wang, X. J. The pro-inflammatory role of TGFβ1: A paradox? Int. J. Biol. Sci. 8, 228–235 (2012).

36. Nagaev, I., Bokarewa, M., Tarkowski, A. & Smith, U. Human resistin is a systemic immune-derived proinflammatory cytokine targeting both leukocytes and adipocytes. PLoS One 1, e31 (2006).

37. Li, M. O. & Flavell, R. A. TGF-β: A master of all T cell trades. Cell 134, 392–404 (2008).

38. Shaw, P. J. & Feske, S. Regulation of lymphocyte function by ORAI and STIM proteins in infection and autoimmunity. J. Physiol. 590, 4157–4167 (2012).

39. Shevach, E. M. Mechanisms of Foxp3+ T Regulatory Cell-Mediated Suppression. Immunity 30, 636–645 (2009).

40. Zhang, G. & Ghosh, S. Negative regulation of toll-like receptor-mediated signaling by Tollip. J. Biol. Chem. 277, 7059–7065 (2002).

41. Blanco, J. C. G. et al. Interferon regulatory factor (IRF)-1 and IRF-2 regulate interferon γ-dependent cyclooxygenase 2 expression. J. Exp. Med. 191, 2131–2144 (2000).

42. Jefferies, C. A. Regulating IRFs in IFN driven disease. Front. Immunol. 10, (2019).

43. Coomes, S. M. et al. CD4 + Th2 cells are directly regulated by IL-10 during allergic airway inflammation. Mucosal Immunol. 10, 150–161 (2017).

44. Couper, K. N., Blount, D. G. & Riley, E. M. IL-10: The Master Regulator of Immunity to Infection. J. Immunol. 180, 5771–5777 (2008).

45. Xu, S. & Cao, X. Interleukin-17 and its expanding biological functions. Cell. Mol. Immunol. 7, 164–174 (2010).

46. Li, C., Corraliza, I. & Langhorne, J. A defect in interleukin-10 leads to enhanced malarial disease in Plasmodium chabaudi chabaudii infection in mice. Infect. Immun. 67, 4435–4442 (1999).

47. Gazzinelli, R. et al. In the Absence of Endogenous IL-10, Mice Acutely Infected with Toxoplasma gondiii Succumb to a Lethal immune Response Dependent on CD4+ T Cells and Accompanied by Overproduction of 11-12, IFN-y, and TNF-cw. J. Immunol. 157, 798–805 (1996).

48. Gu, Y. et al. Interleukin 10 suppresses Th17 cytokines secreted by macrophages and T cells. Eur. J. Immunol. 38, 1807–1813 (2008).

49. Kulcsar, K. A., Baxter, V. K., Greene, I. P. & Griffin, D. E. Interleukin 10 modulation of pathogenic Th17 cells during fatal alphavirus encephalomyelitis. Proc. Natl. Acad. Sci. U. S. A. 111, 16053–16058 (2014).

50. Taylor, C. H. et al. Physiological, but not fitness, effects of two interacting haemoparasitic infections in a wild rodent. Int. J. Parasitol. 48, 463–471 (2018).

51. Dinarello, C. A. Historical insights into cytokines. Eur. J. Immunol. 37, 34–45 (2007).

52. ter Horst, R. et al. Host and Environmental Factors Influencing Individual Human Cytokine Responses. Cell 167, 1111-1124.e13 (2016).

53. Li, Y. et al. A Functional Genomics Approach to Understand Variation in Cytokine Production in Humans. Cell 167, 1099-1110.e14 (2016).

54. Salnikova, L. E. et al. Cytokines mapping for tissue-specific expression, eQTLs and GWAS traits. Sci. Rep. 10, 1–14 (2020).

55. Turner, A. K., Begon, M., Jackson, J. a., Bradley, J. E. & Paterson, S. Genetic Diversity in Cytokines Associated with Immune Variation and Resistance to Multiple Pathogens in a Natural Rodent Population. PLoS Genet. 7, e1002343 (2011).

56. Viney, M. & Riley, E. M. The immunology of wild rodents: Current status and future prospects. Front. Immunol. 8, 1–9 (2017).

