## Supplementary Information for "Early-life immune expression profiles predict later life health and fitness in a wild rodent"

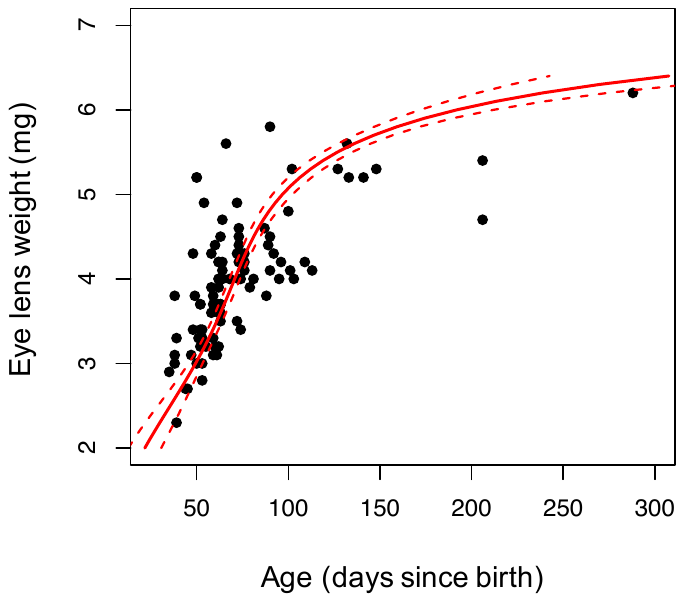

**Fig. S1** Relationship between age and eye lens weight for those voles first sampled prior to breeding (for which we had an approximate birth date) that were culled in later life as part of our cross-sectional study (*n* = 99). Relationship modelled using quassipoisson GLM with quadratic and cubic terms for eye lens weight, and log link. Solid line indicates prediction from the model; red dashed lines indicate 95% confidence intervals.

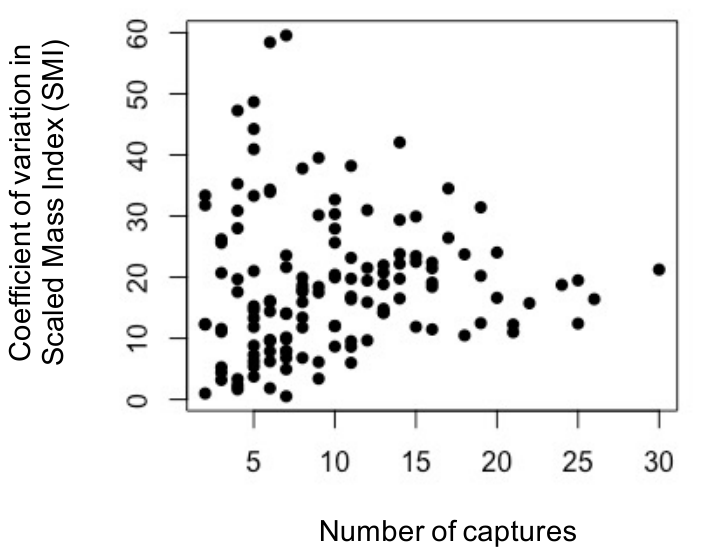

**Fig. S2** Number of times an individual was captured against coefficient of variation in Scaled Mass Index (SMI; a measure of an individual’s average condition across their lifetime^1^). There is no obvious trend but, as expected, values appear to be more variable (and presumably less reliable) when based on fewer captures.

**Supplementary References**

1.	Peig, J. & Green, A. J. New perspectives for estimating body condition from mass/length data: The scaled mass index as an alternative method. *Oikos* **118**, 1883–1891 (2009).
